# Cesium-137 and X-Ray Irradiation Yield Comparable Immune Phenotypes and Activation States in Bone Marrow Chimeric Studies

**DOI:** 10.64898/2026.08.25.746967

**Authors:** Alexander G. Bastian, Eric W. Livingston, Matthew P. Zimmerman, Amy G. Reynolds, Wan Lin Chong, Emily K. Cox, Haiyang Wang, Hong Yuan, Brian C. Miller

**Affiliations:** Lineberger Comprehensive Cancer Center, University of North Carolina at Chapel Hill, Chapel Hill, North Carolina, USA; Biomedical Research Imaging Center, University of North Carolina at Chapel Hill, Chapel Hill, North Carolina, USA; Department of Cell Biology and Physiology, University of North Carolina at Chapel Hill, Chapel Hill, North Carolina, USA; Department of Microbiology and Immunology, University of North Carolina at Chapel Hill, Chapel Hill, North Carolina, USA; Department of Pharmacology, University of North Carolina at Chapel Hill, Chapel Hill, North Carolina, USA; Department of Radiology, University of North Carolina at Chapel Hill, Chapel Hill, North Carolina, USA; Department of Medicine, Division of Oncology, University of North Carolina at Chapel Hill, Chapel Hill, North Carolina, USA; Department of Genetics, University of North Carolina at Chapel Hill, Chapel Hill, North Carolina, USA

## Abstract

Bone marrow chimeras are widely used to study immune development and function. As the field moves from cesium-137 (^137^Cs)-based irradiators to X-ray irradiators for safety reasons, it is essential to determine if there are differences in immune system reconstitution after irradiating mice with one of these two radiation sources. Here, we performed a comprehensive immunological comparison of mice lethally irradiated with ^137^Cs or one of two different X-ray platforms and reconstituted with congenic bone marrow. Mice received 12 Gy total body radiation in two 6 Gy sessions followed by intravenous transfer of donor hematopoietic stem cells and were analyzed eight weeks post-transplant. We assessed mouse survival, donor chimerism, immune cell subset distribution, and activation states across multiple organs (bone marrow, spleen, lymph nodes, liver, and lung). All groups exhibited comparable survival and high levels of donor chimerism, with expected organ-specific reconstitution patterns. Immune lineage distributions, CD4/CD8 ratios, and activation states did not differ by irradiation type. Host-derived radioresistant cells were also similar across all irradiation groups and were predominantly composed of T cells skewed toward an activated phenotype. Overall, our data show that X-ray irradiation with proper filters and energy levels (225 KVp and 320 KVp) can yield equivalent immunological outcomes, including immune reconstitution and activation states, as compared to the same radiation dose from ^137^Cs-based irradiation in bone marrow chimera models. These results support the continued adoption of X-ray irradiation systems in place of ^137^Cs for generating bone marrow chimeras to be used across a wide range of immunologic studies.

## Introduction

Bone marrow chimeras are a vital tool in immunological research allowing genetic manipulation of stem cells and their daughter cells as well as lineage tracing (1–3). Bone marrow chimeras are achieved by lethal irradiation of the host to deplete the endogenous hematopoietic system followed by adoptive transfer of donor hematopoietic stem cells (HSCs). Radiation-induced clearance creates open niches within the bone marrow, spleen, and other immunological tissues, facilitating engraftment of transplanted HSCs. These donor cells reconstitute the immune system, giving rise to all cell lineages. The efficiency of immune reconstitution varies by lineage and tissue, with myeloid cells typically recovering more rapidly than lymphocytes, which depend on selection and peripheral expansion (12–14). Bone marrow chimeras have been used to understand the ontogeny and plasticity of macrophages, the requirements for thymic selection and peripheral tolerance in T cells, the dynamics of B cell development and antibody responses, as well as other foundational discoveries in the mechanisms underlying stem cell development and hematopoiesis (4–8). Moreover, this model is critical for understanding tissue-specific immune reconstitution following irradiation and the interplay between hematopoietic and non-hematopoietic cells in disease models ranging from cancer to infection to autoimmunity (9–11). As such, bone marrow chimeras remain a powerful and versatile system for investigating fundamental and translational questions in immunology.

Ionizing radiation exerts its biological effects by depositing energy into tissues and producing ionization events. These events can directly damage DNA or generate secondary electrons that induce radiolysis of water, producing reactive oxygen species that drive DNA damage, including double-strand breaks (15–17). Hematopoietic stem cells are highly susceptible to radiation-induced DNA damage due to their proliferative nature and limited DNA repair capacity compared to quiescent, radioresistant stromal cells (18). Lethal doses of total body irradiation (typical LD_50/30_ is 7–10 Gy depending on mouse strain, sex, age, radiation source, etc.) induce widespread depletion of hematopoietic cells, including bone marrow progenitors, peripheral lymphocytes, and most tissue-resident immune populations. The resultant pancytopenia is fatal within 10–14 days post-irradiation unless rescued by bone marrow transplantation (19–21).

While lethal irradiation is highly effective at depleting the immune compartment, across murine models roughly 5-15% of immune cells remain following irradiation, with the precise fraction varying by tissue (12,22). The surviving cells are disproportionately composed of memory T cells, which represent the dominant radioresistant immune subset (23,24). The relative radioresistance of T cells has been attributed to several, non-mutually exclusive mechanisms including enhanced DNA repair mechanisms (25,26), cell cycle quiescence (27), and increased anti-apoptotic capacity (28,29). Still, the vast majority of memory T cells are cleared following whole-body irradiation, as evidenced by the fact that patients require re-vaccination after stem cell transplant.

Cesium-137 (^137^Cs) irradiators have been the gold standard for bone marrow chimera generation due to their availability and relatively simple protocols (30,31). These machines generate high energy gamma photons through a source of radioactive ^137^Cs. ^137^Cs (t_1/2_=∼30 years) naturally undergoes beta decay to ^137m^Ba, which then emits gamma photons at ∼662 keV energy (31). However, the use of ^137^Cs irradiators poses significant safety and security risks due to the potential for radiological contamination, environmental hazards, and misuse as a source for ‘dirty bombs,’ prompting a global shift toward safer, non-radioisotope alternatives (32). Clinically, most cesium-137 or cobalt-60 irradiators have been replaced with X-ray irradiators, which generate X-rays by colliding accelerated electrons into a metal target, although research scientists have been more reluctant to switch (33). One reason for researchers’ concern is that whereas X-ray irradiators produce radiation with variable and broad energy levels depending on the power configuration of the machine (35,36), photons generated in ^137^Cs based irradiators are within a narrow energy window which peaks at 662 KeV. The different energy level and spectrum profiles between the two radiation sources can result in different tissue penetration and absorption, thus potential differences in radiobiological effects. In addition to mitigating the safety concerns, transitioning research models to X-ray irradiators may improve the translatability of pre-clinical animal studies, given that patients now receive predominantly X-ray irradiation (34). Although a few studies have compared these irradiation sources for bone marrow chimera studies, the results of these studies have been varied.

One of the earliest systematic comparisons in murine models was conducted by Gibson *et al*., who evaluated lethal-dose and hematopoietic reconstitution following ^137^Cs or X-ray irradiation prior to bone marrow transplantation (BMT) (30). The peak energy of the X-ray source used in that study was 160 KVp. Their study showed significant differences in LD_50/30_ and BMT reconstitution between the two sources, raising concern about their equivalence for BMT experiments. Gott *et al.* evaluated bone marrow ablation and splenocyte ablation by either ^137^Cs or X-ray (320 KVp) irradiation at three different doses. The study concluded that 320 kVp X-ray radiation was suitable for bone marrow ablation, but less effective for splenocyte depletion than ^137^Cs radiation (22). In their study, the corresponding X-ray dose was 30% less than the dose from ^137^Cs radiation (assuming X-ray relative biological effectiveness (RBE) of 1.3, based on their prior modeling studies), suggesting that a higher X-ray dose may be needed to achieve equivalency, especially for splenocyte depletion. Several subsequent studies reported comparable bone marrow chimerism from ^137^Cs and 350 KVp X-ray sources at similar radiation doses in tumor immunology and autoimmunity studies (37,38). However, these studies mainly focused on immune reconstitution in blood or spleen. The chimerism in non-lymphoid organs and immune activation states have not been fully elucidated in studies comparing ^137^Cs and X-ray irradiation. Additionally, many studies lack careful irradiator calibration and dose verification.

In this study, we tested the reconstitution of non-lymphoid and lymphoid organs, comparing Cesium and two X-ray irradiators with different energy levels (225 KVp and 320 KVp) at the same radiation dose (6Gy administered twice) as used in ^137^Cs radiation. We performed a deep immunological characterization of immune cells and their activation states across multiple tissues (spleen, lymph node, bone marrow, liver, and lung). We found no differences in immune reconstitution nor immune activation states between Cesium and X-ray irradiators across any organ, with at least 85% donor reconstitution in all tissues. The remaining host cells were predominantly T cells which had an activated/memory CD44+ phenotype. Overall, our study suggests that labs using bone marrow chimeras for immunologic studies can transition from ^137^Cs to X-ray irradiators and achieve equivalent immune reconstitution throughout the recipient tissues, although X-ray energy level, dose, and filters must be carefully considered.

## Materials and Methods

### Animal models

CD45.1 donor (C57BL/6J-*Ptprc^em6Lutzy^*/J) and CD45.2 recipient (C57BL/6J) congenic female mice aged 6–8 weeks were used for all experiments. Mice were bred and maintained in the University of North Carolina at Chapel Hill Department of Comparative Medicine animal facility. Prior to irradiation, all animals were housed under specific pathogen-free (SPF) conditions in a barrier facility. All experimental groups were maintained under identical housing conditions. After radiation, all mice were housed in autoclaved, individually ventilated cages and provided autoclaved rodent chow (LabDiet 5V0F) and water *ad libitum*. Animals were maintained on a 12-hour light/dark cycle under controlled temperature and humidity conditions. These studies were approved by the University of North Carolina Institutional Animal Care and Use Committee.

### Irradiation

The radiation exposures were conducted using three irradiator systems managed by the Research Radiation Core (RRC) facility: a cesium-137 based irradiator (Gammacell-40 Exactor model, Best Theratronics, Ltd), a standard X-ray irradiator (X-Rad320 model, Precision X-ray, Inc), and an image-guided X-ray irradiator (SmART+ model, Precision X-ray, Inc).

The Gammacell-40 irradiator produces gamma rays generated by dual cesium-137 sources. The system was calibrated using the Fricke dosimetry method by Best Theratronics (the irradiator vendor) every two years. The most recent calibration results are shown in Supplemental Table 1. The Fricke dosimetry showed that the dose variation between the center and edge was 2.4%, and the variation between the measured and the expected dose was less than 2.5%. Variation in the entire chamber (260 mm diameter and 100 mm height) is 7% according to the manufacturer’s specifications. Radiation dose rate at the time of animal radiation for the cesium irradiator was 0.77 Gy/min.

The X-Rad320 system uses the GE ISOVOLT-320 TITAN X-ray unit (4000 Watts output power) with oil-to-air cooling to generate X-ray with maximum energy at 320 KVp for radiation treatment. The irradiator was calibrated using a Farmer type ion chamber (PTW TN30010, Freiburg, Germany) by R3 X-Ray LLC (Hudson, FL) before the comparison study following AAPM TG-61 protocol for absolute dosimetry (39). The calibration was performed at a source to detector distance of 50 cm. Radiochromic film dosimetry was conducted to verify the radiation dose for each radiation study. Detailed radiochromic film dosimetry is provided in Supplemental Table 1. For this study, animals were placed on the stage with the source-to-subject distance (SSD) of 50 cm. A composite filter with 0.25 mm of copper, 1.5 mm of aluminum, and 0.75 mm of tin was used for whole body radiation. The radiation field was set at 203×203 mm, which is the maximum open field on the SSD-50 stage with the adjustable collimator. Dose variation within the field was assessed by the film dosimetry with 5.8% in the radiation field (detailed diagram in Supplemental Figure 1). A field illuminator is provided to guide the animal placement in the center of the radiation field.

The SmART+ system is a small-animal irradiator with image guidance capability, although image guidance was not used for whole body radiation involved in this study. It uses a Comet power supply and X-ray tube (4500 Watts output power) with water-to-water cooling to generate X-rays with the peak energy level of 225 KVp. The system was calibrated using a Farmer type ion chamber (PTW TN30013 model, Freiburg, Germany) following AAPM TG-61 protocol by the irradiator vendor. Two modifications from the TG-61 protocol were made. A square 4×4 cm field was used instead of the recommended 10×10 cm field. The calibration was performed at a source to detector distance of 30.6 cm rather than the 100 cm recommended. The filter with 0.3 mm copper was used for whole body radiation treatment. The calibrated radiation dose rate was 1.96 Gy/min on the irradiation stage for whole-body radiation with SSD of 564 mm. A small radiation field with 120 mm diameter was defined on the stage to ensure homogenous radiation dose. The dose variation within this circle was measured at 1.6%. The detailed stage setup is shown in Supplemental Figure 2. Table 1 shows the main parameters of each irradiator used in this study.

**Table 1.**
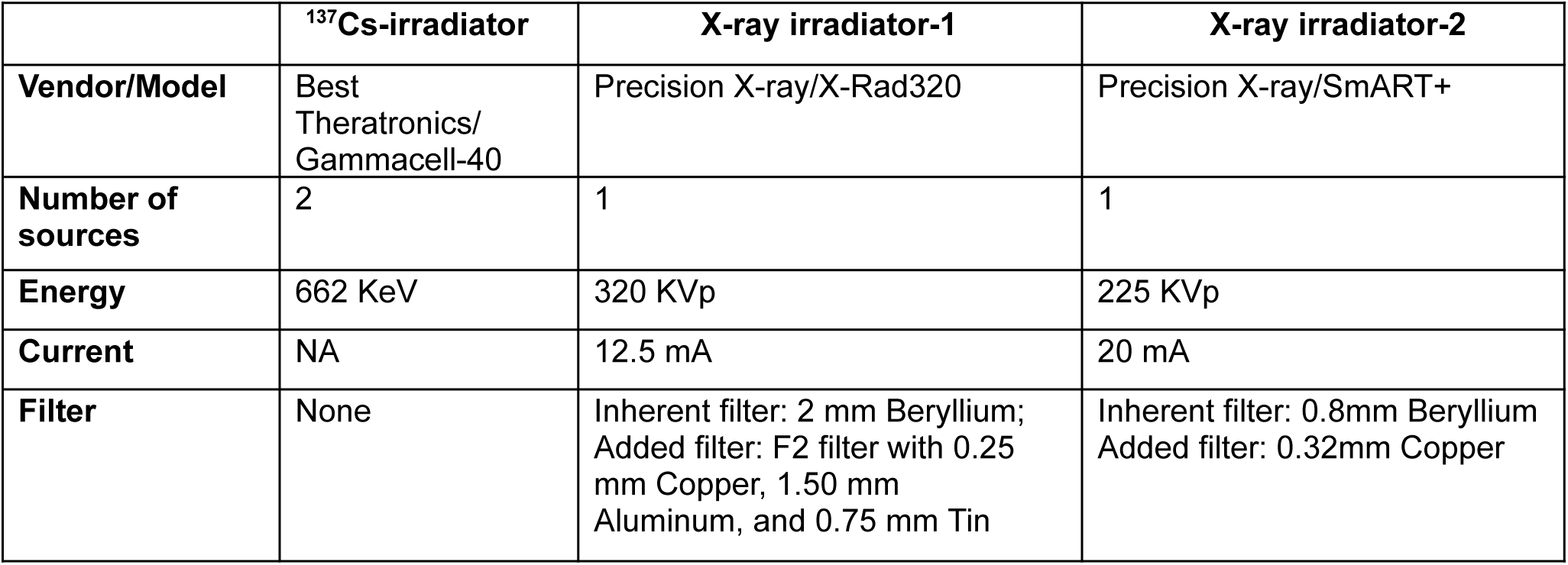
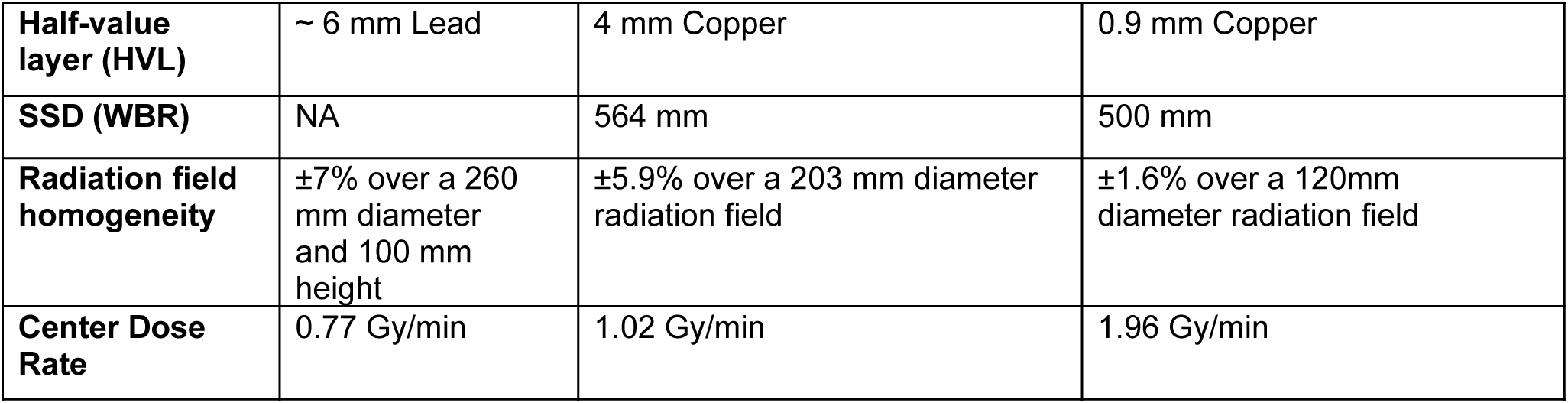
Main parameters of irradiators used in the study.

Filter integration is an important step in X-ray irradiation. Filters are used to remove low-energy (soft) photons, the process usually referred to as “hardening” of the X-ray beam. Filtration can make the dose more uniform in different tissues and ensure enough penetration to deep tissue, such as bone marrow, without causing excessive skin absorption and damage. In addition to the filters inside the X-ray tube (the inherent filters), additional filters are usually applied to shape the beam profile suitable for whole body animal irradiation. The detailed filter information is listed in Table 1. The half-value layer (HVL) value, that is the thickness of material required to reduce the beam intensity by half, is also provided in Table 1 to characterize the beam intensity and quality. As shown there, the X-ray beams from the X-Rad320 system have higher HVL compared to the X-rays from the SmART+ system.

For each experiment, the mice were divided randomly into three groups, and each was assigned to a different irradiator. All groups received two sessions of whole-body radiation with 6 Gy per session, approximately 3 hours apart, for a total radiation dose of 12 Gy. For ^137^Cs irradiation, mice were placed in a circular, perforated, compartmented cage holding 3∼5 mice, and the cage was placed in the middle of the irradiation chamber. For X-ray irradiation, mice were placed in batches of 2-3 mice in a perforated plastic container (Rubbermaid, 8×8×7.5 cm^3^) with a polyethylene lid. The small container size was used to restrain animals from moving during radiation and ensure the dose homogeneity in the radiation field. No anesthesia was used in any radiation procedure, because radiation time per session was short, ranging from 3-7.8 min depending on irradiator type.

### Radiochromic film dosimetry

Radiochromic film dosimetry was performed for X-ray radiation to verify the doses given. A standard calibration curve was first established by exposing the film (Gafchromic EBT3 film, Ashland Inc.) to known doses determined in the treatment planning system in the SmART system based on ion chamber calibration. After 24 hours, the radiochromic films were digitized using an Epson V850 Pro scanner. When scanning, each film was placed in the center region of the scanner bed in a consistent orientation relative to the original sheet. The film images were collected with 16-bit per color channel in three channels at a resolution of 1200 dpi. The films were scanned using a high-performance film scanner (Epson V850Pro) with 1200 dpi resolution. The optical density (OD) of each film was determined based on the ratio to the background film with no exposure. The relationship between the OD and radiation dose was modeled using the following polynomial equation, and the coefficients (*a, b, c*) were determined by a non-linear regression using MATLAB:

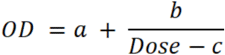

The standard calibration curves for the film batch used are shown in the Supplemental Figure 3. Before or after each X-ray radiation, the radiochromic films from the same batch, as used for standard calibration, were placed between two solid water blocks (VW-3005, CNMC Company, Inc.) with 15 mm of bottom block and 10 mm of top block, which mimics an average mouse size (25 g). The film-block sandwich was placed in the center of the radiation field and underwent the same dose of radiation exposure. The film was processed and scanned following the exact protocol as used in developing standard curves, and the radiation dose was estimated based on the fitted polynomial model.

### Bone Marrow Isolation and Injection

CD45.1 mice were euthanized by CO_2_ asphyxiation and the femurs, tibias, hip bones, and spines were dissected. The bones were cleaned of excess tissue and ground in a mortar and pestle to liberate the bone marrow. The mixture was filtered twice to remove pieces of bone. The bone marrow cells were then mixed with CD117 positive selection beads and sorted by magnetic-associated cell sorting (MACS) to enrich the stem cell population (Miltenyi Biotec). These cells were washed and 500,000 cells per mouse intravenously injected via the tail vein into the irradiated mice 1 hour after the final irradiation. Mice were provided with drinking water containing 2% sulfatrim by volume for 1 week after irradiation.

### Organ processing

8 weeks after irradiation and stem cell transfer, bone marrow chimera mice were euthanized, and the tibias, spleens, livers, lungs, and inguinal lymph nodes dissected. Bone marrow was flushed from the femurs and tibias using a 27G needle and red blood cells were lysed using Ack Lysis Buffer (Gibco). Spleens and lymph nodes were smashed through a 70um filter to prepare a single cell suspension. Red blood cells in the spleens were also lysed using Ack Lysis Buffer. The livers and lungs were chopped into small pieces (approximately 2 mm in diameter) and incubated at 37C for 45 minutes in a digestion buffer containing 50ug/mL DNase (Sigma-Alrich) and 1mg/mL Collagenase 1 (Millipore-Sigma). The cells were then passed through a 70um filter to create a single cell suspension. Cells isolated from livers and lungs were then incubated with CD45 positive selection beads (Miltenyi Biotec) and sorted by MACS to isolate the CD45+ immune cells.

### Flow Cytometry

Cells were incubated with anti-CD16/32 antibody cocktail (Fc Block) (Biolegend) and mouse serum to block Fc receptors for 30 minutes. Cells were then incubated with either an extracellular lymphoid or myeloid master mix of pre-diluted antibodies (Supplemental Table 2). After 30 min incubation at 4C, cells were fixed with IC fixation buffer (eBiosciences) for 30 minutes at 4C then permeabilized with Permeabilization Buffer (eBiosciences) for 30 minutes containing Fc block (Biolegend). Anti-Foxp3 was then added for intracellular staining in the lymphoid panel. Single stain controls were prepared using UltraComp compensation beads. Flow cytometric analysis was performed on a Symphony A3 and data analyzed using FlowJo software. The sequential flow gating strategy is shown in Supplemental Figure 4.

### Blinding Procedure

All tissue processing, flow cytometric staining, data acquisition, and analysis were performed by an investigator blinded to the irradiation group assignments. Sample identifiers were coded by a separate team member responsible for irradiation and animal handling, and decoding occurred only after data analysis was complete. This approach minimized potential bias in gating strategies, cell subset quantification, and interpretation of immunological differences between irradiation types.

### Statistics

All statistical analyses were performed using GraphPad Prism and/or Microsoft Excel. Comparisons between irradiation groups within each organ were conducted using unpaired two-tailed Student’s *t*-tests. For each immune population and activation parameter, pairwise comparisons were performed independently within each tissue. To account for the large number of comparisons across tissues and immune subsets, *p*-values were adjusted using the Benjamini–Hochberg false discovery rate (FDR) correction. This approach controls the expected proportion of false positives while maintaining statistical power in high-dimensional datasets. A threshold of 0.05 was considered statistically significant. In cases where unadjusted statistical significance was observed but did not persist after FDR correction, results were interpreted as not statistically significant.

## Results

To directly test the impacts of Cesium versus X-ray irradiation on immune cell depletion and reconstitution, mice were irradiated in a cesium irradiator or with one of two different X-ray irradiators with 6 Gy administered twice (3 hours apart) for a total of 12Gy. Radiochromic film dosimetry was conducted at least three times before or after the X-ray irradiation to validate the dose. With 6 Gy prescribed radiation dose, the average dose measured from the film dosimetry was 6.19 ± 0.23 Gy (mean ± SD) for the X-Rad320 system, and 5.86 ± 0.06 Gy for the SmART system.

The animals then received 500,000 hematopoietic stem cells from a congenically labeled donor (Figure 1A). There was no difference in mouse survival between the irradiation methods: 79% (X-Rad320-Xray), 86% (Smart+-Xray)), and 93% survival (Gammacell-40-Cs) (Figure 1B). Survival percentage for all groups was comparable with our lab’s overall average survival in chimeric studies using Cesium (82% survival from 54 total mice over 3 independent experiments). The mice that did not survive all died the first 2 weeks after radiation, likely due to unsuccessful immune reconstitution. After 8 weeks, the bone morrow, inguinal lymph nodes, spleens, lungs and livers of all mice were harvested. We observed expected differences in the presence of radioresistant cells across organs; lowest in bone marrow (2% host cells), highest in the lymph nodes (16% host cells). However, we did not identify any differences between irradiation groups (Figure 1C).

**Figure 1:**
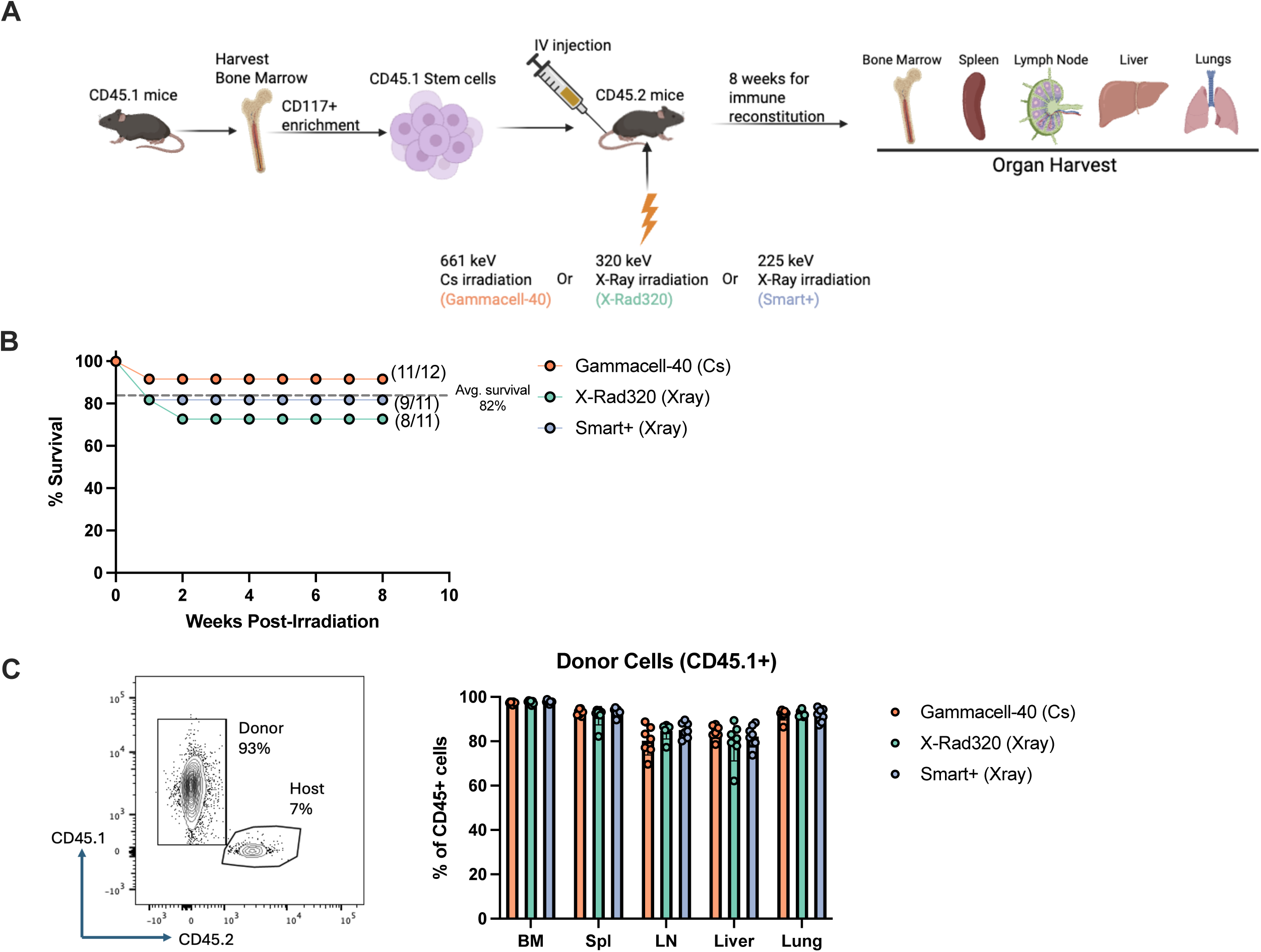
No difference between Cesium-137 or X-Ray irradiation was observed in survival or chimerism. (A) Experimental design comparing the effects of X-Ray and ^137^Cs irradiation to generate bone marrow chimeras. CD45.2 mice were irradiated twice with 6 Gy, 3 hours apart. Mice were then IV injected with 500,000 stem cells from CD45.1 mice for congenic tracking of donor and host cells. After 8 weeks, the bone marrow, spleen, inguinal lymph nodes, liver, and lungs were harvested and immune cells analyzed by flow cytometry. (B) Survival curves after irradiation and HSC reconstitution. Average survival shown from past chimera experiments using Cs irradiation. (C) After 8 weeks, chimerism was assessed by CD45.1% of total CD45+ cells across all organs. Representative flow plots from the spleen shown. Data shown from one of two replicate experiments.

Given prior reports of differences in specific immune population reconstitution between Cesium and X-ray irradiation (30), we next analyzed immune subsets between the radiation groups. Differences in immune makeup were observed between different organs, with the lymph nodes showing a strong skewing towards the lymphoid lineage (T cells and B cells comprising 95%) and the bone marrow showing a much larger myeloid skewing (50%). We observed no differences between radiation groups in the percentages of reconstituted T cells, B cells, granulocytes, or non-granulocyte myeloid cells (Figure 2A-C).

**Figure 2:**
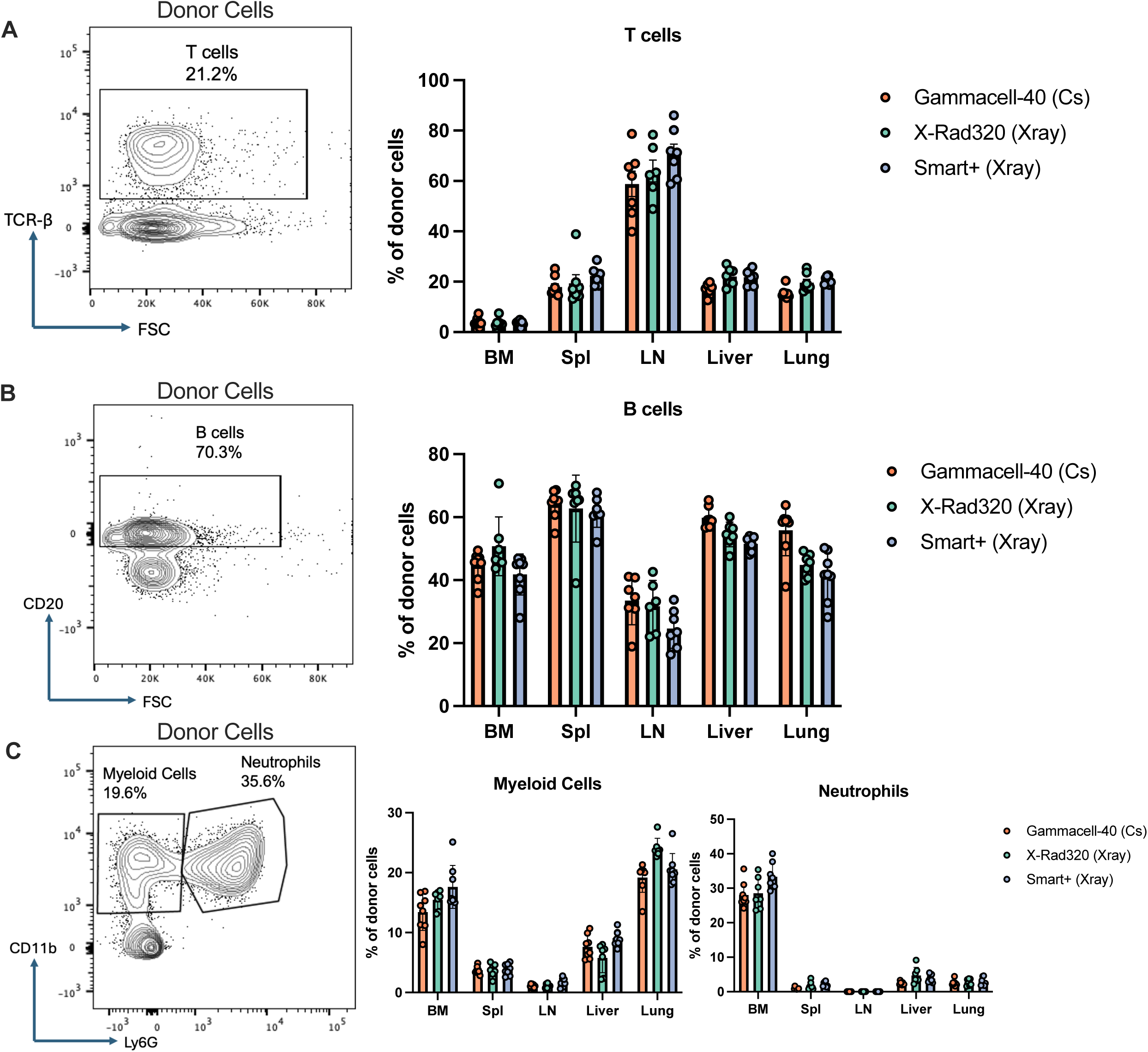
Global lymphoid and myeloid compartment reconstitution were unaffected by radiation type. Gated on live CD45.1+ donor cells, percentages of (A) T cells (TCR-β+) (B) B cells (CD20+) and (C) myeloid cells (CD11b+Ly6G-) and neutrophils (CD11b+Ly6G+) are shown. Representative flow plots from the spleen shown for (A) and (B). Representative flow plot from the bone marrow is shown in (C). Data shown from one of two replicate experiments.

T cells are key effectors of pathogen control, autoimmunity, and anti-tumor immunity. To determine if Cesium and X-ray irradiation would yield equivalent T cell reconstitution for downstream studies, we analyzed T cell subsets and activation states across various tissues, excluding bone marrow due to the low T cell abundance. No significant differences in CD4^+^/CD8^+^ T cell ratios (2.5:1 in all radiation groups in the spleen) were detected between radiation groups (Figure 3A). Similarly, the proportion of immunosuppressive Foxp3^+^ regulatory T cells remained within 5–10% across all tissues and was unaffected by irradiation type (Figure 3B). Next, we measured activation states of the different T cell populations as measured by CD44 and CD62L expression. No significant differences were found in activation states of reconstituted CD4^+^ T cells (Figure 3C). The percentages of CD44^+^CD62L^+^ memory T cells were similar across radiation groups (ranging between 38% and 41% between radiation groups in the spleen, for example). Similarly, CD44^-^CD62L^+^ naïve T cell numbers were comparable across radiation groups (ranging between 42-43% of splenic CD4+ T cells) as were CD44^+^CD62L^-^ effector T cells (13-15%) (Figure 3C). Similarly, no differences were observed in reconstituted CD8^+^ T cells when analyzing memory, naïve and effector T cells across all tissues (Figure 3D). Therefore, our data indicate that reconstituted T cell subset distribution and activation state are unaffected by irradiation method.

**Figure 3:**
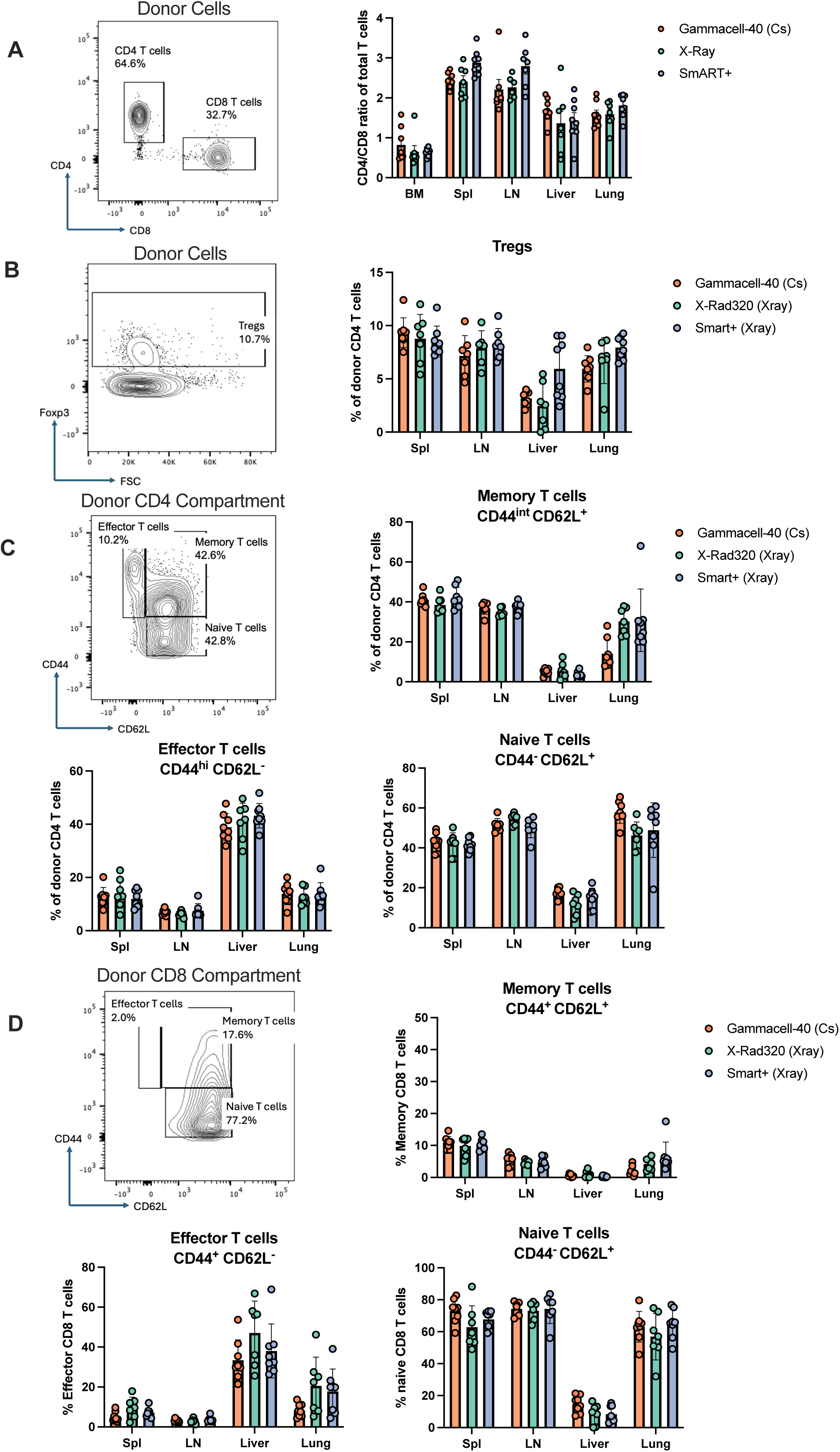
Cesium-137 and X-Ray irradiation led to indistinguishable donor T cell phenotypes and activation states. (A) Ratio of CD4+ to CD8+ T cells in harvested organs. (B) Proportions of Treg cells identified by intracellular Foxp3 expression in total CD4+ T cells. (C-D) Activation states of CD4+ T cells (C) or CD8+ T cells (D) assessed by CD44 and CD62L expression. Representative flow plots from the spleen shown. Data shown from one of two replicate experiments.

Next, we analyzed reconstitution and activation of non-neutrophil myeloid cells (CD11b+Ly6G-) after bone marrow transplantation. No differences between radiation groups were observed within tissues when analyzing the percentage of monocytes (Ly6C^+^F4/80^-^). In the spleen, the percentage of monocytes within the myeloid compartment ranged from 37% to 41%. Similarly, we observed no difference in macrophage reconstitution in all organs (Figure 4A). Within the spleen, we observed a range of 21-24% macrophages (F4/80^+^Ly6C^-^) within the myeloid compartment (Figure 4A). We also analyzed the activation status of monocytes and macrophages based on MHC II expression across tissues, excluding the lymph node due to the minimal myeloid abundance in that compartment. We found no significant differences between the activation status of monocytes (Figure 4B) nor macrophages (Figure 4C) based on radiation type received. Additionally, no differences were observed in the lung for alveolar macrophages (SiglecF^+^CD64^+^) between radiation groups (Figure 4D). Collectively, our results demonstrate that cesium and X-ray irradiation result in equivalent myeloid cell reconstitution.

**Figure 4:**
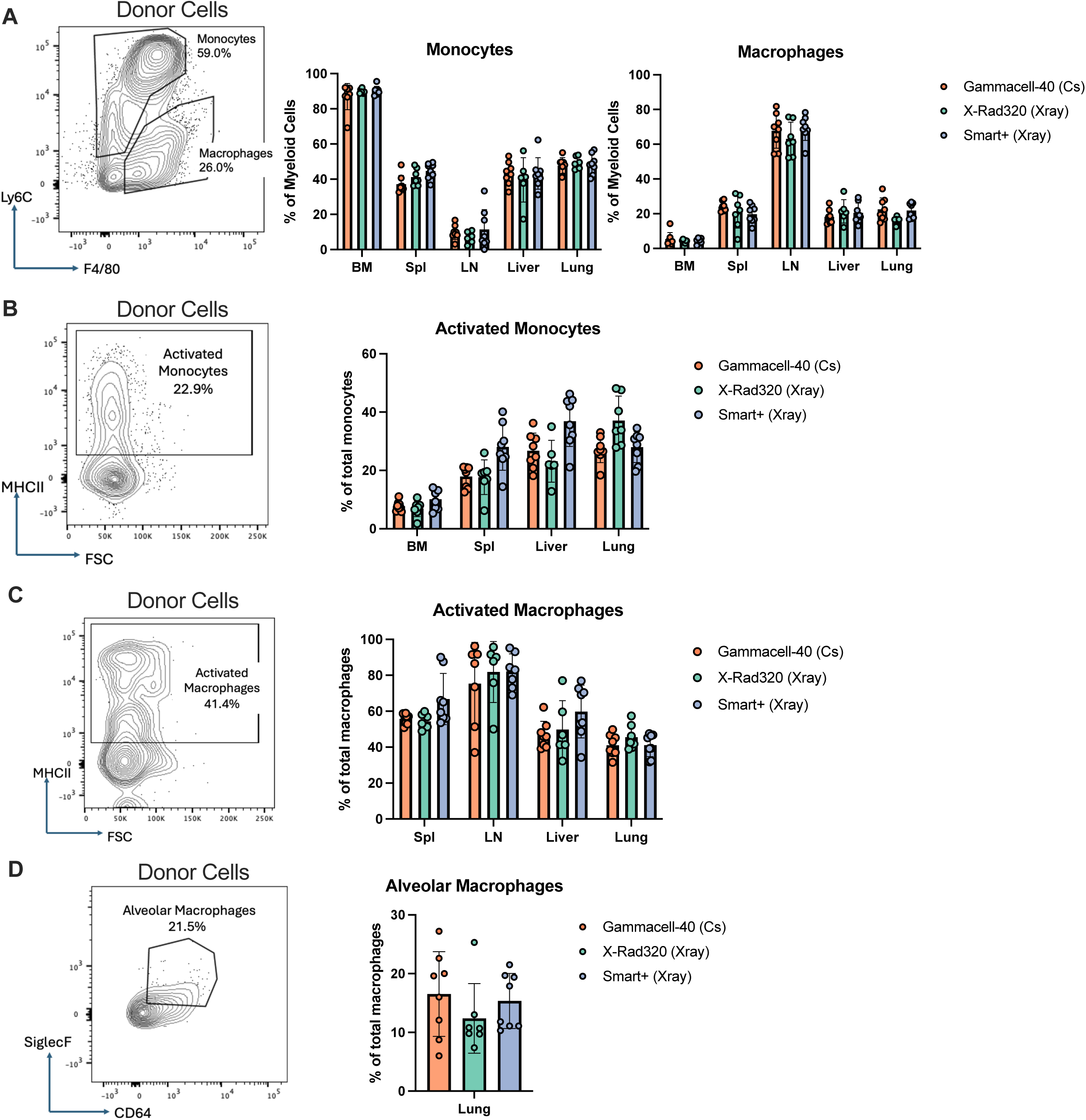
Cesium-137 and X-ray irradiation led to comparable myeloid cell reconstitution and activation. (A) Proportion of myeloid cell populations as a total of all non-neutrophil myeloid cells (CD11b+Ly6G-). Monocytes (Ly6C+F4/80-) and macrophages (Ly6C-F4/80+) quantified. (B-C) Activation is determined by surface expression of MHC II on the monocyte (B) or macrophage populations (C). Quantification of alveolar macrophages, defined by SiglecF and CD64 co-expression within the Ly6C-/F4-80+ macrophage gate from lung. Representative flow plots shown from the lung. Data shown from one of two replicate experiments.

Finally, we analyzed the surviving host cells (CD45.2^+^) to identify if there are differences in remaining immune cells after cesium or X-ray irradiation. As previously shown (Figure 1C), there were no differences observed between radiation groups in terms of the chimerism seen in all organs. Independent of the radiation group or tissue type, the majority of radioresistant cells were T cells (83-93% across tissues) (Figure 5A). We observed no differences in the CD4+/CD8+ T cell ratios or Treg percentage (Supplemental Figure 5A-B) in the host cell compartment between radiation groups. In the host CD4+ T cell compartment, no difference was observed in the activation state of the cells based on effector (CD44+CD62L-), memory (CD44+CD62L+), and naive (CD44-CD62L+) classifications (Figure 5B). Similarly, no difference among the activation subsets was observed within the host CD8+ T cell compartment (Figure 5C). When comparing the donor and host T cell compartments, a clear trend emerged across all tissues and radiation groups: the host compartment was skewed towards an activated T cell phenotype (CD44+, expressed on effector and memory cells). Our data show that activated T cells are more radioresistant compared to naïve T cells, with these cells surviving for many weeks after radiation exposure. This skewing was observed in both the CD8+ (Figure 5D) and CD4+ T cell compartments (Supplemental Figure 5C).

**Figure 5:**
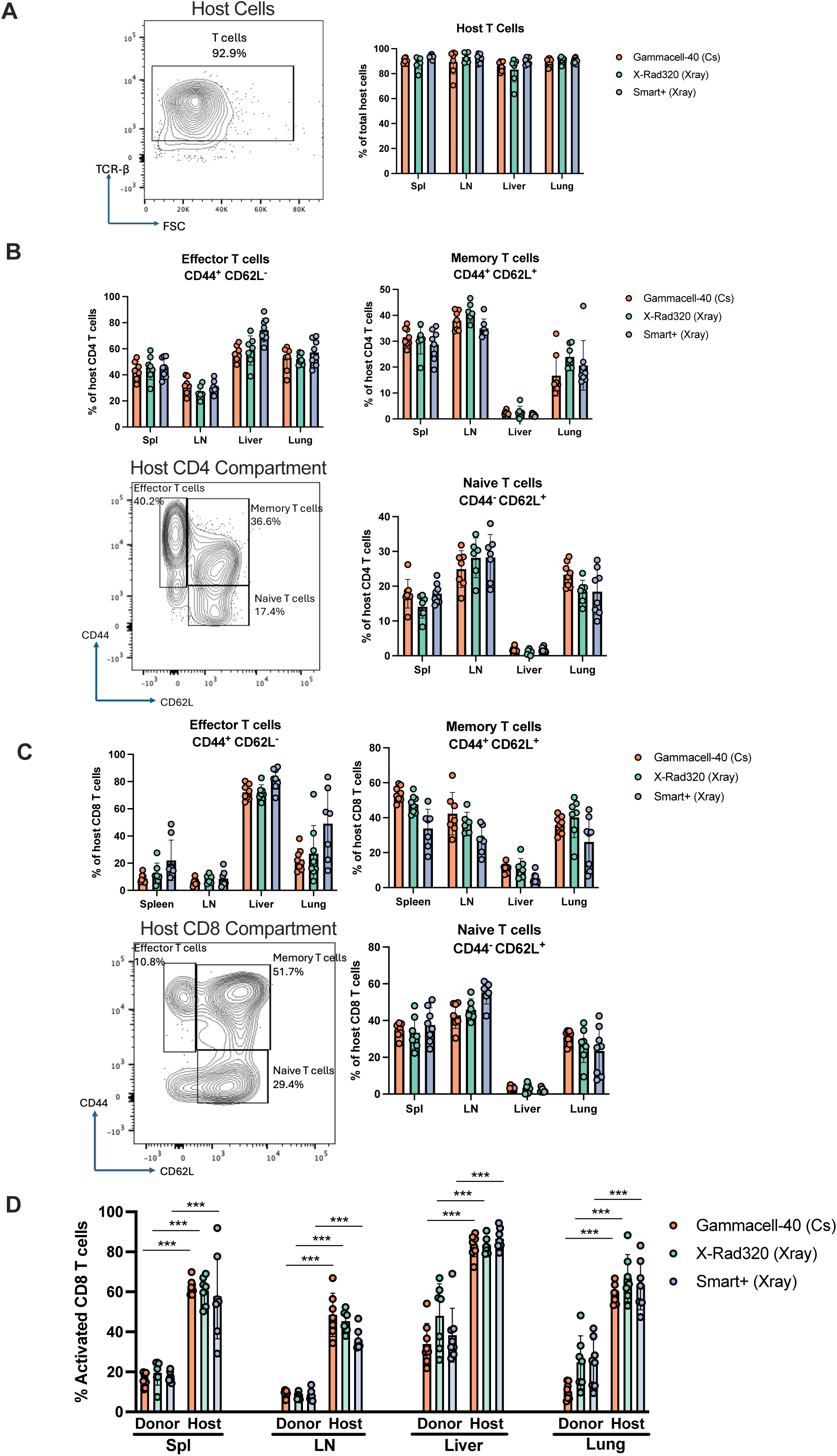
Activated T cells made up the majority of the radioresistant cells in both Cesium-137 and X-ray irradiated mice. (A) Percentage of TCR-β+ T cells of total host (CD45.2+) cells shown. (B-C) T cells were stratified by CD4 and CD8 expression. Of the CD4+ T cells (B) and CD8+ T cells (C), the activation state of the cells is determined by CD44 and CD62L expression. (D) Percentage of CD44+ (combination of effector and memory) CD8+ T cells comparing host and donor across all organs. All representative flow plots shown are from the spleen. Data shown from one of two replicate experiments.

## Discussion

Taken together, these results provide comprehensive evidence that immune reconstitution is equivalent in both lymphoid and non-lymphoid tissues after appropriately dosed Cesium or X-ray irradiation. No meaningful differences between radiation methods were observed in donor-derived immune profiles or the radioresistant host immune cells. Specifically, we observed no differences in mouse survival, donor engraftment, tissue-specific reconstitution, immune subset frequencies, and cell activation states in our study between irradiation platforms. These results strongly support the conclusion that X-ray irradiation serves as a suitable and biologically equivalent alternative to Cesium irradiation for bone marrow chimera generation. Furthermore, by incorporating two independent X-ray platforms (225 KVp and 320 KVp), our study demonstrates that the equivalence between ^137^Cs and X-ray irradiation extends across different machine configurations. These findings support the ongoing push towards replacing Cs-based irradiators with safer X-ray irradiators.

Our results complement and extend prior studies comparing ^137^Cs and X-ray irradiation in bone marrow transplantation and chimera models. As mentioned earlier, Gibson *et al.* reported that although ^137^Cs and X-ray methods were both capable of supporting donor cell engraftment, there were significant differences in lineage-specific immune reconstitution depending on the radiation source (30). It should be noted that the peak energy level used for X-ray irradiation in the Gibson *et al.* study was 160 kVp, which is lower than that used in our study (225 KVp and 320 KVp). In addition, it was not clear what type of filter was used in this study, which could significantly affect the X-ray spectrum profile and tissue absorption. Studies have shown that low energy 160 KVp X-rays have much less tissue penetration and greater dose differences between bone and soft tissue, compared to the higher energy 320 KVp X-rays or gamma photons from ^137^Cs (31), potentially resulting in insufficient radiation to the bone marrow and higher skin irritation. The two X-ray irradiators used in this study provided different X-ray qualities with different peak energy (225 KVp vs. 320 KVp) and HVL (0.9 mm copper vs. 4 mm copper, respectively). However, no significant differences in immune reconstitution or activation states were observed between the two X-ray irradiation groups. Based on our study, we believe that integrating high X-ray energy (225 KVp or up) and a beam-hardening filter could result in comparable radiation effects with ^137^Cs irradiation.

Another question is what an equivalent dose is when switching from ^137^Cs to X-ray radiation, i.e., what an appropriate RBE value is. The photon absorption and thus radiobiological effects in tissue is energy dependent (31,40). With decreased energy, there is more tissue absorption and thus increased hematopoietic and intestinal injury. It is suggested that reduced X-ray dose may be used to achieve comparable biological effects (40). Prior work has suggested RBE ranging from 1.1 to 1.3 when transitioning from ^137^Cs to X-ray radiation (22,37,38). Gott *et al.* showed that 320 KVp X-ray at 30% reduced dose compared with ^137^Cs radiation produced less effective splenocyte ablation, suggesting that X-ray RBE of 1.3 may be overestimated - a higher X-ray dose might be needed (22). Wittenborn *et al.* showed 11 Gy of X-ray radiation (RBE of 1.15) produced comparable hematopoietic chimerism as compared to the 13 Gy of ^137^Cs irradiation and described that 13 Gy of X-ray caused much higher mortality, thus BMT chimera could not be compared effectively (38). In many of these studies it is unclear exactly how radiation was administered, as detailed radiation procedures and dose calibration were not provided. In our study, where radiation was administered in two split sessions with 6 Gy per session, we have shown that the equal dose of X-ray (RBE=1) produced comparable survival and immune reconstitution as compared to the same dose of ^137^Cs radiation.

Dose consistency across X-ray platforms is also an important consideration. We noticed that the radiation dose was more variable from the X-Rad320 system, ranging from 5.85 Gy to 6.44 Gy for a prescribed 6Gy dose. In contrast, the SmART X-ray system provided a more consistent radiation dose ranging from 5.76 to 5.93 with about 2% variation from the prescribed dose. The X-Rad320 system is a much older X-ray system, purchased in 2010, while the SmART system was installed in 2023. The X-ray radiation was also limited in that there was no turning table in either of the systems to provide a more homogenous radiation field. We used a small animal holding box to minimize radiation variation in one field. We also used the field illuminator in both systems to guide the placement of animal cages.

It has been shown that dose rate plays an important role in the biological effects from radiation (31,41). In our study, the ^137^Cs irradiator provided the lowest dose rate at 0.77 Gy/min, followed by X-Rad320 at 1.02 Gy/min, and SmART system at 1.96 Gy/min. The differences in dose rates may not be dramatic with the range between ∼0.5-2 Gy/min. Thus, variation in dose rate in our study likely does not contribute significantly to the overall radiobiological responses.

Importantly, residual host-derived immune populations, representing the radioresistant fraction, were predominantly T cells, with no shifts in identity or abundance between radiation groups. We noticed the greatest retention of these cells in the lymph nodes and livers of host mice, highlighting the importance of considering tissue-specific immune environments when interpreting host–donor dynamics in bone marrow chimera studies. Notably, these residual host T cells consistently displayed a more activated phenotype (CD44+) relative to the donor-derived T cell population across all tissues. This observation aligns with prior evidence suggesting that antigen-experienced or memory T cells exhibit enhanced radioresistance, likely due to differences in cell cycle status or upregulated survival pathways (24,25,27,28,42). The persistence of these cells up to eight weeks after lethal irradiation, with their conserved activation profile, suggests that radioresistance of antigen-experienced T cells is independent of the radiation source.

By performing a comprehensive analysis of immune composition and activation state across multiple immunologically relevant organs, we provide a higher-resolution assessment of immune reconstitution following irradiation. These tissues represent distinct immunological environments that differ in immune composition, activation status, and exposure to antigens, making them particularly informative for evaluating the fidelity of immune recovery after bone marrow transplantation. Importantly, assessing immune activation states provides functional context beyond lineage distribution, as the activation status of immune cells strongly influences their behavior in downstream experimental disease models such as infection, cancer, and autoimmune disease. While the transition from Cesium to X-ray irradiators should be approached thoughtfully, particularly for groups with long-standing protocols, our data strongly support the use of X-ray irradiation as comparable to Cesium for bone chimera generation without compromising immunological fidelity.

## Author Contributions

Conception and design: AGB, EWL, HY, BCM

Acquisition of data: AGB, EWL, MPZ, AGR, WLC, EKC, HW

Analysis and interpretation of data: AGB, EWL, HY, BCM

Writing, review, and/or revision of the manuscript: AGB, EWL, HY, BCM

Study supervision: HY, BCM

## Acknowledgements

Radiation was conducted in the UNC Research Radiation Core (RRC) Facility. Flow cytometry was conducted in the UNC Flow Cytometry Core (FCC) Facility (RRID:SCR_019170). Colony management services were performed by the Colony Management Core within the Division of Comparative Medicine as well as the UNC Lineberger Preclinical Research Unit (PRU). The RRC, FCC, and PRU are supported in part by the NCI Center Core Support Grant (P30CA016086) to the UNC Lineberger Comprehensive Cancer Center (LCCC). Aaron F Gunsalus at the UNC Radiation Safety Office assisted with the operation of the cesium-137 irradiator and the X-RAD320 X-ray irradiator.

## Supplemental Figure Legends

**Supplemental Figure 1: Radiation stage and field homogeneity of the X-Rad320 X-ray irradiator.** (A) Radiation chamber and stage setup. There is a field illuminator to provide the radiation field information. The maximum radiation field with the adjustable collimator at the SSD-50cm position is 203 x 203 mm. (B) Measured radiation dose rate at the center and the edge of the radiation field. The coefficient of dose variation in the radiation field is 5.9%.

**Supplemental Figure 2: Radiation stage and field homogeneity of the SmART X-ray irradiator.** (A) Radiation chamber and stage setup for whole body radiation (WBR). A plastic board (9 mm thick) is placed on the top of the metal stage to reduce the backscatter during WBR. Radiation field is marked on the board as a circle. Animal box is placed within the circle to ensure field homogeneity and position consistency. (B) Radiochromic film taken on the stage with an open field setting (no collimator). The circle has a diameter of 120 mm, corresponding to the radiation field marked on the stage. (C) Cross-line profile of the radiation field. The square region corresponds to the range within the circle. The coefficient of dose variation within the circle is 1.6%.

**Supplemental Figure 3: Standard calibration curves for the EBT-3 radiochromic film**. Films (Batch#11292202) were scanned using the Epson V850Pro with a 16-bit dynamic range, as shown in the upper panel. The relationship between radiation dose and film intensity in the red, green, and blue channel was fitted to a polynomial model. The red channel was used to estimate the dose based on film intensity.

**Supplemental Figure 4: Flow gating strategy**. All cells were gated based on FSC/SSC, then single cells and live cells. Cells were then split into donor or host cells; all downstream gating was applied to both.

**Supplemental Figure 5: Radioresistant T cells had a more activated phenotype.** (A) The CD4/CD8 T cell ratio among the CD45.2+ host T cells is shown. (B) The percentage of CD4 T cells that are Foxp3+ are shown in each organ. (C) The percentage of CD44+ cells within the CD4+ T cell compartment is shown. No bone marrow data is shown due to paucity of T cells in the bone marrow.

**Supplemental Table 1: Fricke dosimetry report of the Gammacell-40.** (Serial #:265)

**Supplemental Table 2: Details of Flow Cytometry Antibodies**

