## Supplemental Figures and Tables for "Cesium-137 and X-Ray Irradiation Yield Comparable Immune Phenotypes and Activation States in Bone Marrow Chimeric Studies"

**Supplemental Figure 1**

A

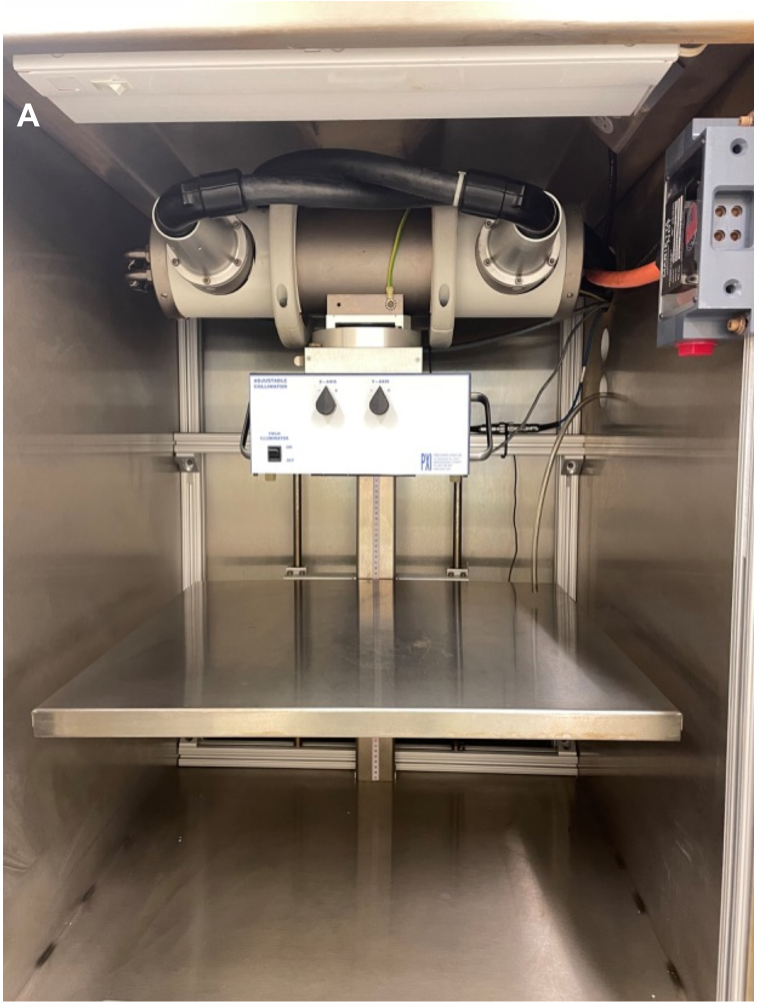

B

| Maximum Radiation Field with adjustable collimator: 203x203 mm | |
|---|---|
| Position | Dose Rate (Gy/min) |
| Center | 1.02 |
| 1 | 0.98 |
| 2 | 0.89 |
| 3 | 0.90 |
| 4 | 0.99 |

### Supplemental Figure 2

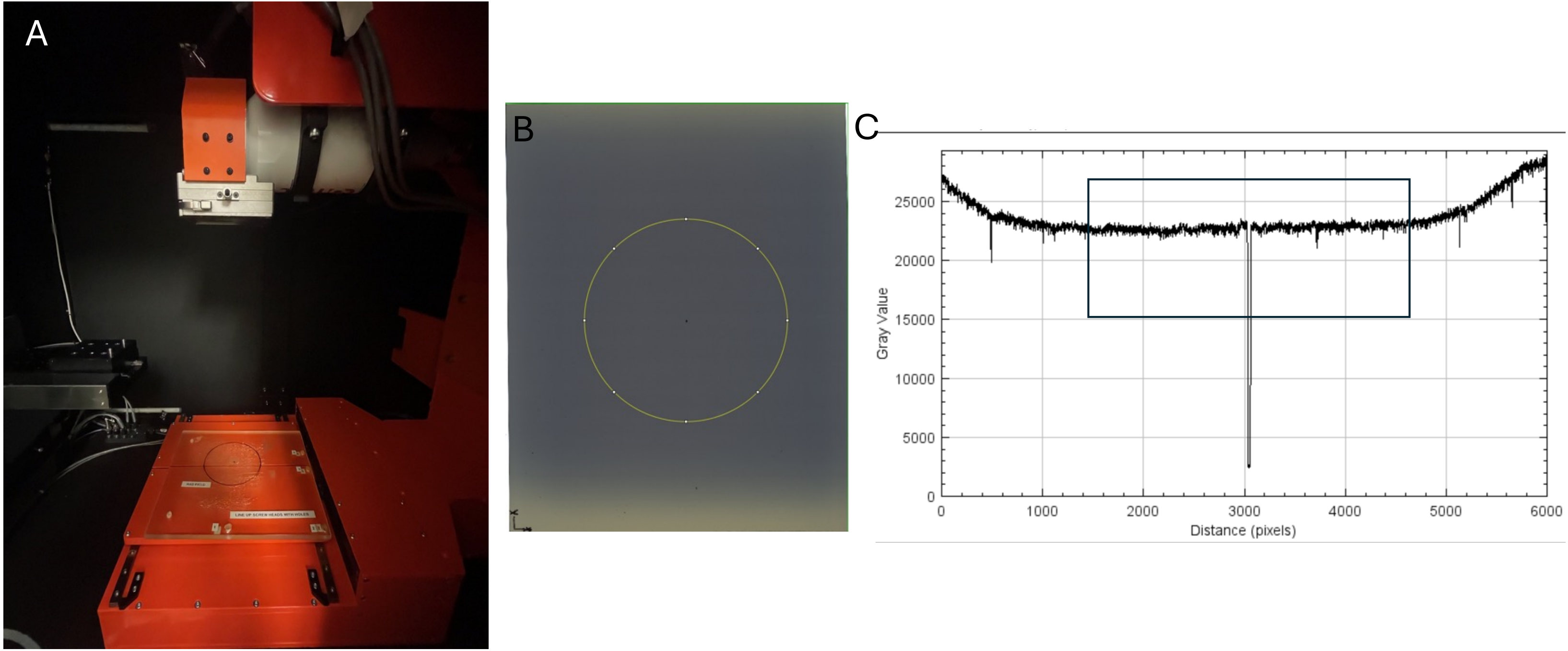

**Supplemental Figure 3**

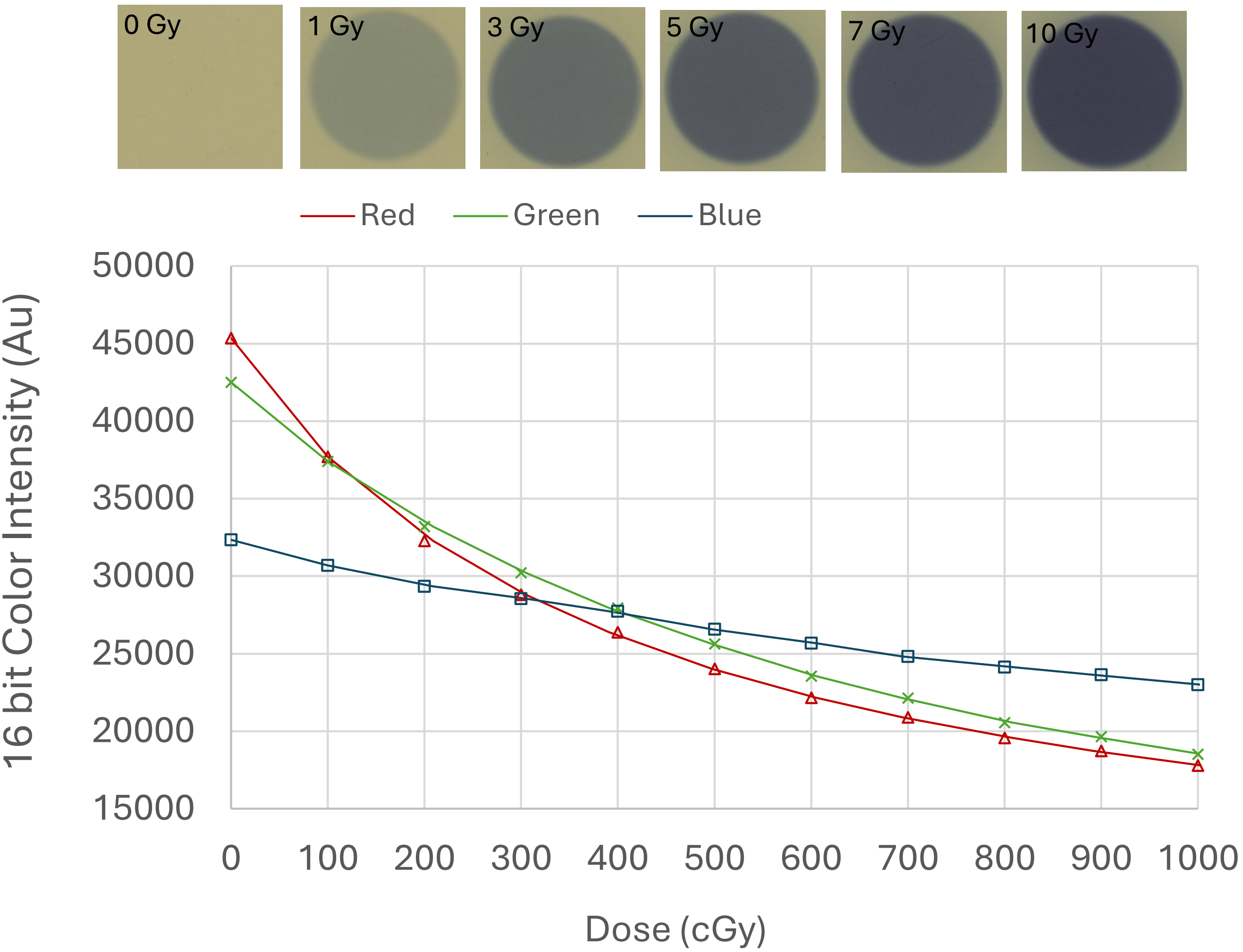

### Supplemental Figure 4

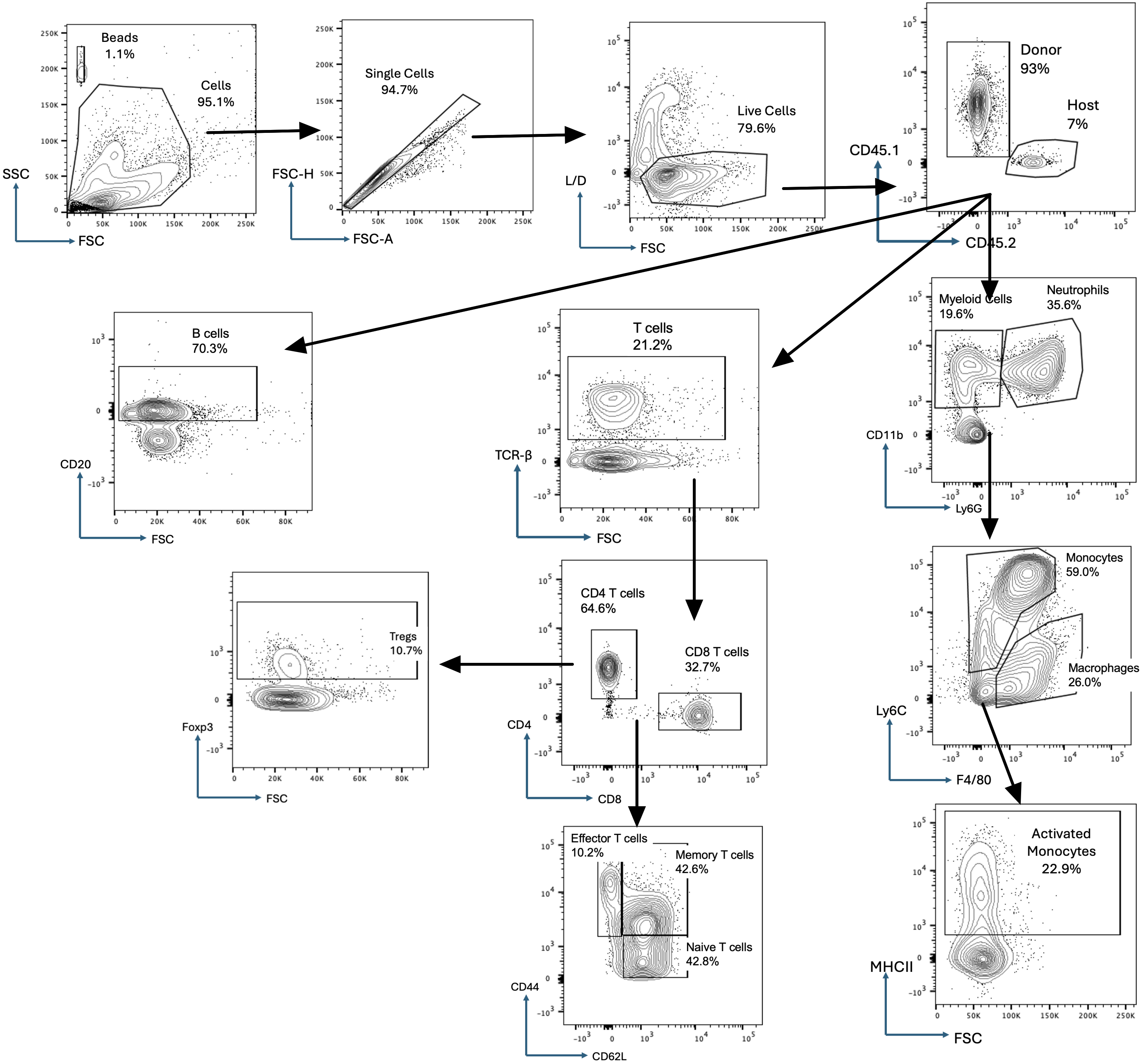

### Supplemental Figure 5

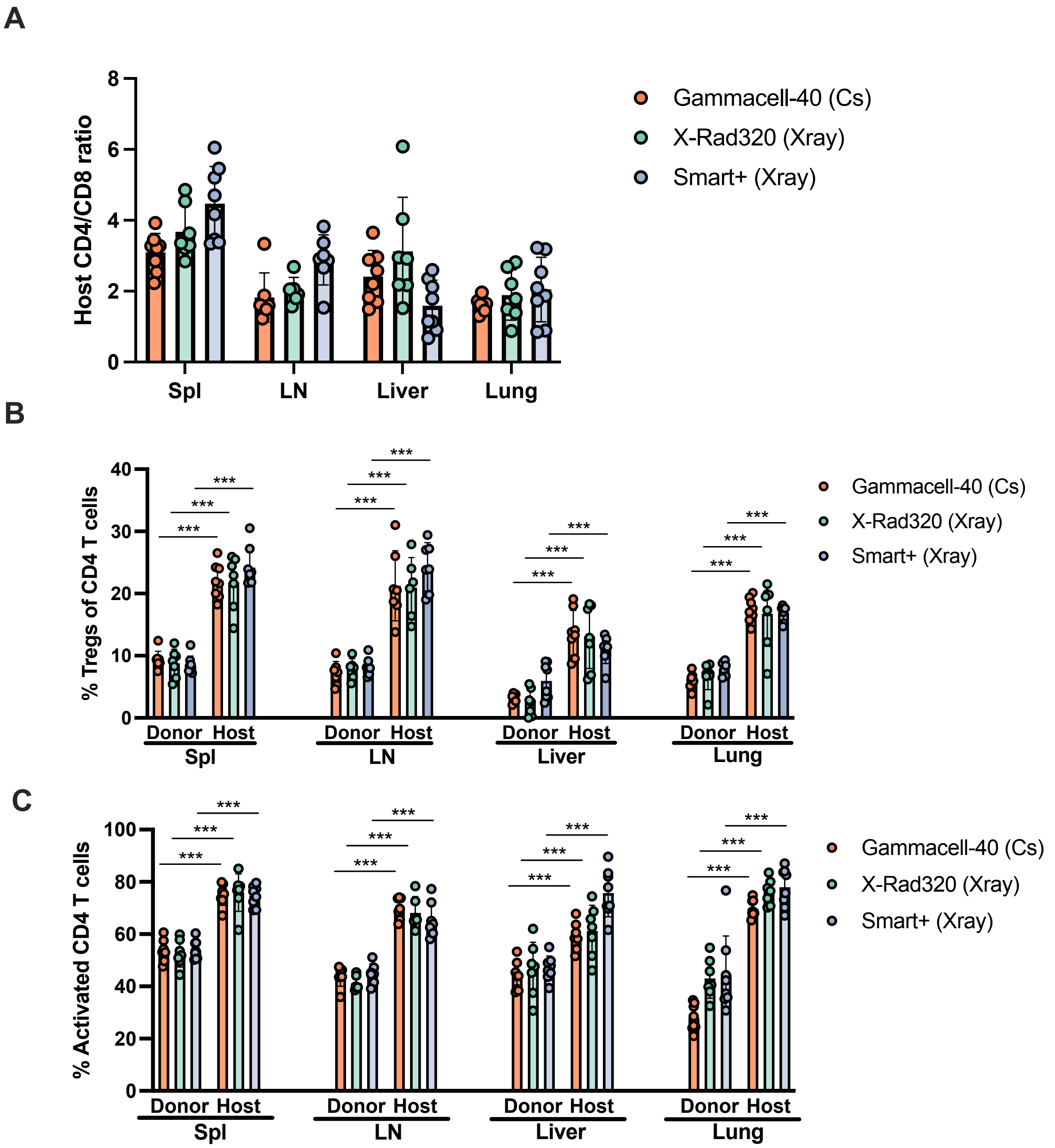

**Supplemental Table 1**

| Position | Measured Absorbed Dose Rate (Gy/min) | Expected Absorbed Dose Rate* (Gy/min) | Percent Difference (Measured to Expected Dose Rate |
|---|---|---|---|
| Central Dose Rate | 0.794 ± 2.5% | 0.812 | -2.2% |
| 3 o'clock position | 0.791 ± 2.5% | 0.794 | -0.4% |
| 7 o'clock position | 0.813 ± 2.5% | 0.824 | -1.3% |

* The expected dose was calculated from the original Best Theratronics measurement done on 4/10/2014, measured at 1 Gy/min.

**Supplemental Table 2**

| Lymphoid Panel | | | | | |
|---|---|---|---|---|---|
| **Marker** | **Fluorophore** | **Clone** | **Company** | **Catalog #** | **Dilution** |
| Live/Dead NIR | - | - | Invitrogen | L34994A | 1000 |
| CD45.1 | PE/Cy7 | A20 | Biolegend | 110730 | 200 |
| CD45.2 | BV605 | 104 | Biolegend | 109841 | 200 |
| TCR-β | BV785 | H57-597 | Biolegend | 109249 | 200 |
| CD4 | PerCP/Cy5.5 | GK1.5 | Biolegend | 100434 | 200 |
| CD8a | PE/Dazzle | 53-6.7 | Biolegend | 100761 | 200 |
| CD44 | AF488 | IM7 | Biolegend | 103016 | 200 |
| CD62L | PE | MEL-14 | Biolegend | 104407 | 200 |
| Foxp3 | APC | FJK-16s | eBiosciences | 17-5773-82 | 100 |
| CD20 | AlexaFluor700 | SA275A11 | Biolegend | 150415 | 200 |

| Myeloid Panel | | | | | |
|---|---|---|---|---|---|
| **Marker** | **Fluorophore** | **Clone** | **Company** | **Catalog #** | **Dilution** |
| Live/Dead NIR | - | - | Invitrogen | L34994A | 1000 |
| CD45.1 | PE/Cy7 | A20 | Biolegend | 110730 | 200 |
| CD45.2 | BV605 | 104 | Biolegend | 109841 | 200 |
| CD11b | BUV396 | M1/70 | BD | 565976 | 800 |
| F4/80 | BUV563 | T45-2342 | BD | 749284 | 800 |
| Ly6C | BUV737 | HK1.4.rMAb | BD | 755201 | 800 |
| CD64 | PerCP/Cy5.5 | X54-5/7.1 | Biolegend | 139308 | 200 |
| SiglecF | BV785 | E50-2440 | Biolegend | 740956 | 100 |
| MHCII | PE | M5/114.15.2 | Biolegend | 107625 | 400 |
| Ly6G | AlexaFluor700 | 1A8 | Biolegend | 127622 | 200 |
